# Lineage plasticity enables low-ER luminal tumors to evolve and gain basal-like traits

**DOI:** 10.1101/2022.11.25.517090

**Authors:** Gadisti Aisha Mohamed, Sundis Mahmood, Nevena B. Ognjenovic, Min Kyung Lee, Owen M. Wilkins, Brock C. Christensen, Kristen E. Muller, Diwakar R. Pattabiraman

## Abstract

Stratifying breast cancer into specific molecular or histological subtypes aids in therapeutic decision-making and predicting outcomes, however, these subtypes may not be as distinct as previously thought. Patients with luminal-like, Estrogen Receptor (ER)- expressing tumors have better prognosis than patients with more aggressive, triple-negative or basal-like tumors. There is, however, a subset of luminal-like tumors that express lower levels of ER, which exhibit more basal-like features. We have found that breast tumors expressing lower levels of ER, traditionally considered to be luminal-like, represent a distinct subset of breast cancer characterized by the emergence of basal-like features. Lineage tracing of low-ER tumors in the MMTV-PyMT mouse mammary tumor model revealed that basal marker expressing cells arose from normal luminal epithelial cells, suggesting that luminal-to-basal plasticity is responsible for the evolution and emergence of basal-like characteristics. This plasticity allows tumor cells to gain a new lumino-basal phenotype, thus leading to intratumoral lumino-basal heterogeneity. Single-cell RNA sequencing revealed SOX10 as a potential driver for this plasticity, which is known among breast tumors to be almost exclusively expressed in Triple Negative Breast Cancer (TNBC) and was also found to be highly expressed in low-ER tumors. These findings suggest that basal-like tumors may result from the evolutionary progression of luminal tumors with low ER expression.

## Introduction

Breast cancer is a complex disease with multiple different biologic subtypes which have clinical implications on tumor development, prognosis, and treatment^1–3^. While traditional surrogate markers can be used to classify breast tumors into hormone-receptor positive, *HER2* amplified, and triple-negative subtypes, advancements in gene expression profiling have helped refine subtype stratification. Gene expression analyses across a diverse range of human breast carcinomas classified these tumors into four intrinsic subtypes: basal-like, *Erb-B2+*, normal-breast-like, and luminal epithelial/ER+^4^. Further refinement of these subtypes based on a larger sample size revealed that the ER-positive luminal epithelial subtype could be further divided into 2 subgroups: luminal A and luminal B, with luminal A expressing higher ER levels than luminal B. These intrinsic subtypes differ in their clinical outcomes, with the basal-like subtype exhibiting the worst prognosis, followed by *Erb-B2+*^5^. A smaller signature of 50 genes, PAM50, may be used by clinicians to classify tumors based on these intrinsic subtypes^6^.

In the clinical environment, immunohistochemistry (IHC) based determination of surrogate protein marker expression is utilized to classify breast carcinomas into four subtypes: 1) Estrogen Receptor (ER) and Progesterone Receptor (PR) positive, and Ki67 low, 2) ER, PR, and Ki67 high, 3) *HER2/neu* amplified, and 4) ER, PR, and *HER2* negative, or triple-negative breast cancer (TNBC). These IHC-based subtypes correspond to the intrinsic subtypes luminal A, luminal B, *Erb-B2+*, and basal-like, respectively, providing biologic and clinically significant information used to guide treatment decisions. Typically, patients with ER negative tumors, TNBC, and *HER2/neu* amplified, benefit from non-hormone-based forms of therapy - adjuvant or neoadjuvant chemotherapy for TNBC tumors, with the addition of anti-HER2 targeted therapy (i.e. Trastuzumab) for *HER2/neu* amplified tumors. Tumors are considered ER-positive when demonstrating 1% or greater ER expression^7^, and patients typically receive treatment with an antiestrogen agent (i.e Tamoxifen) or aromatase inhibitor (i.e. letrozole, anastrozole). This low cutoff for ER-positive determination results in a heterogeneous collection of tumors being considered luminal-like, as tumors with less than 10% ER-positive cells may exhibit different characteristics to those with >10% ER-positivity. There are limited data on the overall benefit of endocrine therapies for patients with low level (1-10%) ER expression, but given the possible benefit, patients are eligible for endocrine treatment^8^. Some studies suggest the majority of breast cancers with low ER expression show molecular features similar to ER-negative, basal-like tumors rather than ER-positive, endocrine sensitive tumors^9^. It is essential to understand the underlying biology of breast cancer with low ER expression, in order to recognize their prognostic significance and identify ideal treatment regimens.

The normal mammary epithelium consists of cells from two different lineages: a luminal lineage characterized by the expression of Keratin 8 (Krt8), with more committed cells expressing ER and PR, and a basal lineage expressing Keratin 5 (Krt5) and/or Keratin 14 (Krt14)^10, 11^. These lineages are derived from a bipotent mammary stem cell (MaSC) progenitor in the embryonic stage, but are maintained postnatally by unipotent luminal and basal progenitors^12–15^. Despite this lineage restriction, several studies have revealed the potential for lineage plasticity in the adult mammary gland in non-homeostatic settings. For example, lineage plasticity of the luminal and basal compartment allows them to regain multipotency in the adult mammary gland with luminal-derived basal cells (LdBCs) emerging in response to hormone stimulation during pregnancy^16^, and basal cells repopulating mammary epithelium in response to injury or luminal cell ablation^17^. In the neoplastic setting, the luminal lineage has been identified as the cell of origin for *BRCA1*-mutant basal-like breast cancers suggesting its involvement in the development of TNBC-like tumors typically observed in these patients^18^. Moreover, *BRCA1* and p53 deletions in the mouse luminal compartment results in tumors resembling typical human basal-like tumors^19^. In addition, claudin-low breast tumors, a mesenchymal subset of TNBCs, may also be derived from the luminal lineage^20^. These findings point to lineage plasticity being a core feature in the process of mammary tumorigenesis whereby luminal tumor cells gain the ability to stray from their lineage-of-origin. The heterogeneity of ER expression within luminal-like tumors provides a starting point to study subpopulations within ER-positive tumors that may be more prone to plasticity and the acquisition of basal-like traits.

In this study, we show that luminal tumors with low-ER expression represent a distinct subtype with a higher tendency to gain basal-like traits. These tumors arise from luminal cells undergoing luminal-to-basal plasticity, leading to the emergence of cells that exhibit a lumino-basal phenotype. This plasticity of luminal tumor cells and presence of lumino-basal heterogeneity within breast tumors likely plays a critical role in their overall aggressive traits, especially their ability to progress and gain metastatic propensity.

## Results

### Low-ER breast tumors exhibit distinct basal-like features

Previous studies have observed that invasive breast carcinomas with low-ER expression, in which less than 10% of tumor cells express ER, share more similarities with TNBCs when compared to tumors that harbored more than 10% ER-expressing cells^21^. We analyzed newly diagnosed invasive breast carcinomas from the pathology database at Dartmouth-Hitchcock Medical Center (DHMC) from 2012-2020 (n=2208) and observed 46 (2.1%) that were classified as low-ER tumors containing between 1-10% ER- expressing tumor cells (Fig 1A). Most, (41 out of 46, 89%) were high-grade invasive carcinomas (Fig S1A) with 1-9% ER-expressing tumor cells (Fig 1B). The intensity of ER expression was also reduced in these low-ER tumors, with 93.5% showing moderate or weaker ER staining (Fig 1C and S1B-D). In contrast to high expressing ER tumors which typically also exhibit some degree of progesterone receptor (PR) expression, most low- ER tumors (76%) were PR negative (Fig 1D). The frequency of *HER2*-positivity was as expected, with 24% of cases harboring *HER2/neu* amplification (Fig 1E).

**Figure 1:**
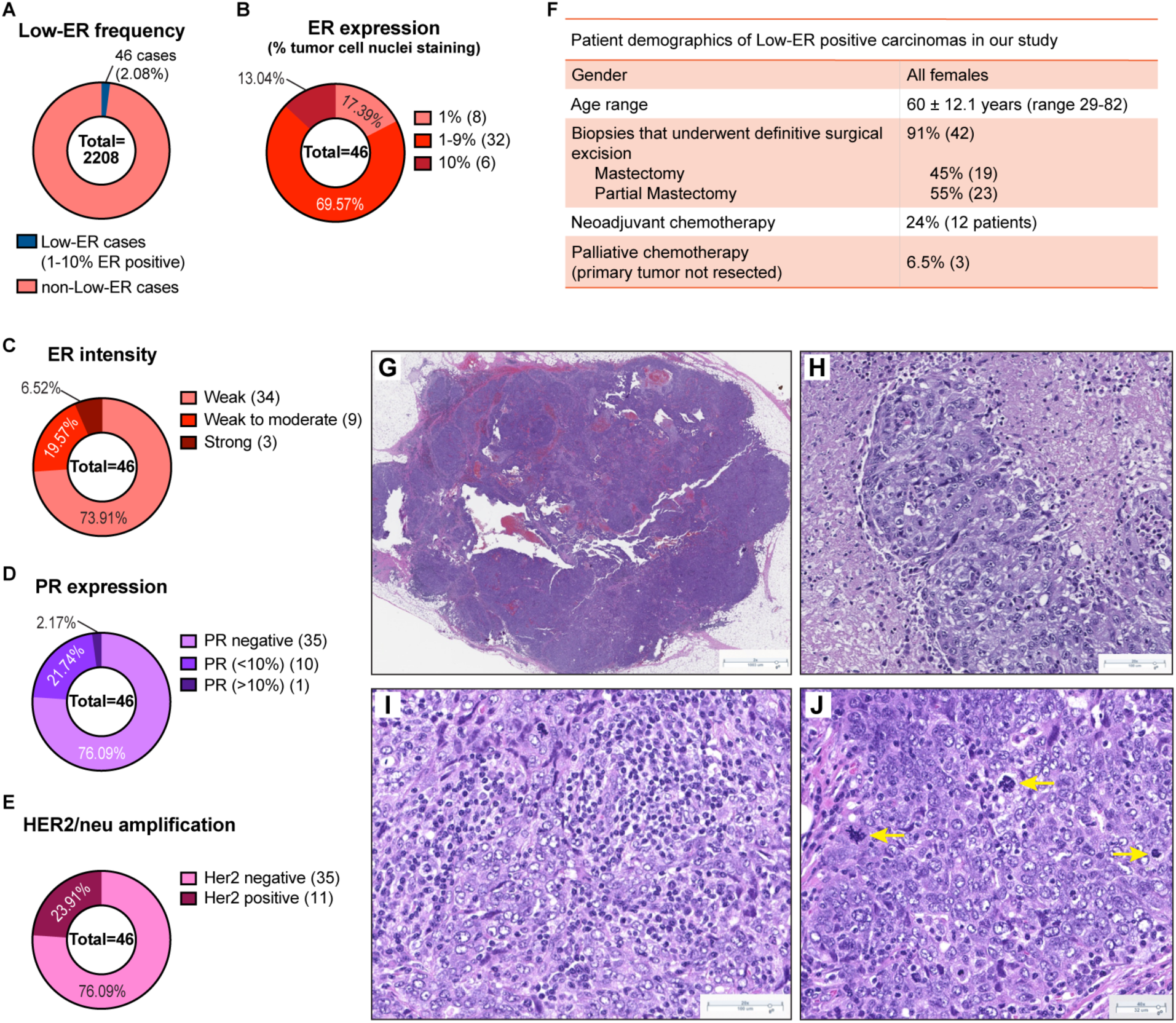
Basal-like features of tumors expressing low ER. **(A)** Frequency of cases expressing low ER levels (<1% ER expressing nuclei) in newly diagnosed invasive breast carcinomas from the pathology database at Dartmouth-Hitchcock Medical Center (DHMC) from 2012-2020. **(B-E)** Breakdown of ER expression levels **(B)**, ER intensity levels **(C)**, PR expression **(D)**, and Her2/neu amplification **(E)**, in the 46 low-ER cases. (**F**) Patient demographics of the 46 low-ER cases in our cohort. **(G-J)** H&E staining of low-ER tumors showing solid tumor with pushing borders **(G)**, pleomorphic, high-grade nuclei with admixed necrosis **(H)**, prominent lymphoplasmacytic infiltrates **(I)**, and conspicuous mitotic activity, including atypical mitoses (yellow arrows) **(J)**.

Treatments administered to patients harboring low-ER tumors are more similar to treatment regimens for patients with TNBC. In our cohort, 43% of patients received chemotherapy, 34% received radiation therapy, and only 23% received hormone therapy (Fig S1E). Interestingly, response rates to neoadjuvant chemotherapy in patients with low-ER tumors were similar to ER-negative tumors and significantly different from tumors with moderate and high ER-positive tumors^22^. twelve patients (24%) received neoadjuvant chemotherapy, and most (75%) achieved a pathologic complete response (Fig 1F and S1A). The majority of patients (81%) had no evidence of the disease at follow-up (Fig S1F).

Microscopic examination of tumors with low-ER expression revealed histologic features commonly present in breast tumors with basal-like molecular profiles and carcinomas harboring *BRCA1* mutations. Twenty of 46 tumors (43%) showed well-circumscribed or pushing borders and were comprised of high-grade, pleomorphic tumor cells arranged in solid sheets with conspicuous mitotic activity, admixed necrosis, and prominent tumor infiltrating lymphoplasmacytic infiltrates (Fig 1G-J). The histologic findings in our cases are in agreement with several other recent studies that have shown that low-ER breast tumors show pathologic characteristics typical of ER-negative tumors with basal-like gene expression profiles^9, 21, 23^.

These data indicate that low-ER tumors are a distinct subtype of breast cancer, separate from the typical, ER-expressing luminal-like subtypes. They display more similarities to basal-like or triple-negative tumors, especially with respect to biomarker expression, pathology and the types of treatments patients receive.

### Low-ER tumors differ from luminal B tumors in their biomarker profiles

To investigate precisely how different low-ER tumors are from luminal tumors, 24 luminal B and 22 low-ER tumors were compiled into two tissue microarrays (TMAs). The luminal B tumors were selected based on a combination of pathologic characteristics including high tumor grade, high mitotic rate (>18 mitoses per 10 high power fields), and diffuse ER expression in tumor cells (all tumors showed >80% tumor cell nuclei with ER expression). Compared to the low-ER group, none of luminal B tumors showed histologic basal-like phenotypic characteristics. In the luminal B group, most patients (87.5%) received hormone therapy, or a combination of hormone therapy and chemotherapy (Fig S2A), which proved effective, with more than 91% of patients showing no evidence of disease at follow-up (Fig S2B). No patients received neoadjuvant chemotherapy (Fig 2A). Most tumors in the luminal B group showed strong PR positivity (Fig S2C) and all tumors were negative for HER2/neu amplification (Fig S2D), which are more typical features of a luminal-like breast cancer subtype. In comparison to low-ER tumors, none of the luminal B tumors contained the constellation of basal-like histologic features we observed in 43% of low-ER tumors.

**Figure 2:**
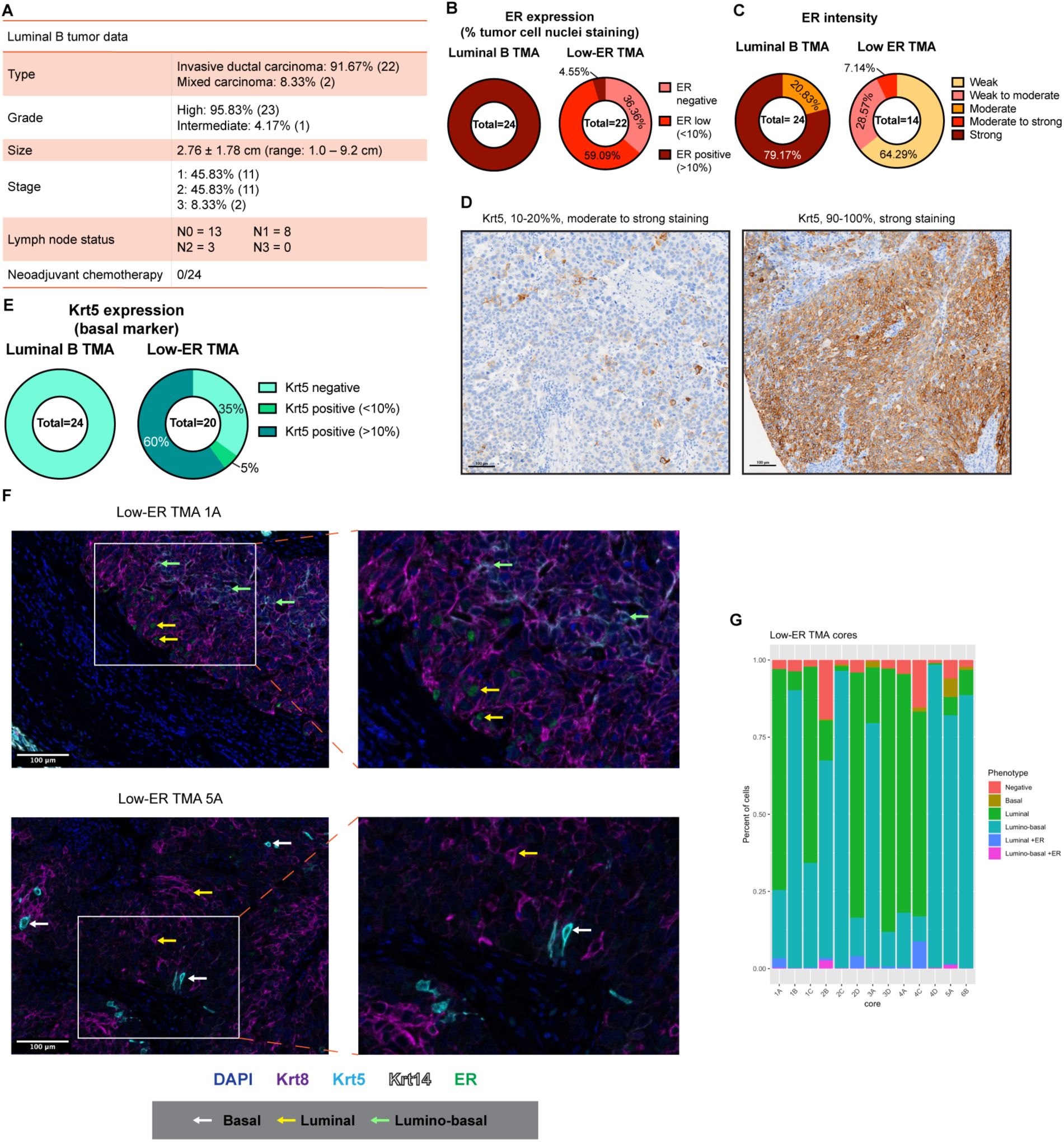
Low-ER tumors are distinct from luminal B tumors. **(A)** Details of the luminal B tumors included in TMAs as a comparison to Low-ER tumors. **(B-C)** ER expression **(B)** and intensity **(C)** levels of the luminal B tumors. **(D)** Representative images of Krt5 IHC staining in luminal B and low-ER TMAs. **(E)** Quantified Krt5 expression in luminal B and low-ER TMAs. **(F)** Representative images of TSA staining containing heterogeneous populations in low-ER TMAs. Samples were stained with Krt8 (purple), Krt5 (cyan), Krt14 (white), ER (green), and DAPI (blue). White arrows point to basal tumor cells, yellow arrows point to luminal tumor cells, and green arrows point to lumino-basal tumor cells. (G) Quantification of different cell phenotypes found within 13 of the low-ER TMA cores which were found to harbor heterogeneous populations (TMA cores with homogenous populations were excluded)

The two TMAs were stained for expression of ER (luminal marker) and Krt5 (basal marker). As expected, immunohistochemistry (IHC) staining of these TMAs showed that all 24 luminal B samples were ER positive (>10% ER expressing cells), with almost 80% showing strong ER intensity, while the low-ER samples were mostly low-ER expressing, with weak or moderate ER intensity (Fig 2B, 2C, and S2E). Eight tumor cores in the low- ER TMA were observed to stain negative for ER, while one had 10-20% ER positive nuclei. The low-ER tumors were identified based on ER expression in the diagnostic biopsy, while TMA cores were obtained from the surgical specimens. These tumors are still biologically low-ER tumors, however due to focal ER expression and heterogeneity, the ER expression of these tumor cores may vary. When stained for Krt5, a basal marker used to identify basal-like breast cancer subtypes, none of the luminal B samples expressed Krt5, but 65% of the low-ER samples were Krt5^+^ (Fig 2D and 2E). Staining for p63, another basal marker, also showed higher positivity in low-ER tumors compared to luminal B (Fig S2F). These data support histological observations that low-ER tumors are more basal-like and express higher levels of basal markers than luminal B tumors.

We wanted to further investigate if the expression of basal markers within the low-ER tumors corresponded to a loss of luminal marker expression. Tyramide Signal Amplification (TSA) staining^24, 25^ was used with luminal marker ER and Krt8 antibodies in addition to basal markers Krt5 and Krt14. We utilized this staining method as it allows the use of multiple antibodies raised in the same species and have previously used it to stain TMAs containing hundreds of patient tumor samples^26, 27^. All luminal B TMA cores strongly expressed both ER and Krt8, with no expression of either basal marker (Fig S2G). In contrast, low-ER TMA cores exhibited more heterogeneous ER, Krt5, and Krt14 expression (Fig 2F). Importantly, the low-ER TMA cores also expressed high Krt8 levels, suggesting luminal lineage identity is retained despite the reduction in ER and increase in Krt5 expression. Along these lines, most of the Krt5-expressing tumor cells also co-expressed Krt8, with only a small percentage of cells exclusively expressing the basal marker.

The low-ER TMA cores were further analyzed to identify the different cell types that these heterogeneous tumors were comprised of. While a few cells were found to only express Krt5, most of the Krt5-expressing cells co-expressed Krt8. Krt14 was less abundant in these tumors, with Krt14^+^ cells also co-expressing both Krt5 and Krt8, indicating that most basal-like cells within these tumors express basal markers without losing their luminal identity i.e., exhibiting a lumino-basal phenotype (Fig 2F). Cells with fully basal phenotypes in which only Krt5 was expressed were rare and only found in 9 out of 13 low-ER tumor cores (Fig 2F and 2G). ER expression was expectedly weak and scarce, but was found both in cells expressing Krt8 only, and in cells co-expressing either Krt8 and Krt5 or Krt8 and Krt14 (Fig 2F).

To analyze the distribution of these various cell types within the low-ER tumors, we quantified each cell phenotype within the low-ER tumor cores. Of the 20 tumor cores analyzed, six cores predominantly consist of the luminal cells, seven cores predominantly consist of cells of the lumino-basal phenotype (Fig 2G), and seven cores were excluded from analysis due to an absence of basal marker expression. As expected, ER expression is more abundant in cells of the luminal phenotype as compared to those exhibiting a lumino-basal phenotype. The strictly basal phenotype was also not commonly found within these tumors, indicating that tumor cells rarely lost all luminal marker expression to become fully basal.

These results provide evidence that distinguishes low-ER tumors from luminal B tumors, both in terms of histopathology and luminal and basal marker expression. Furthermore, low-ER tumors are more heterogeneous in epithelial cell marker expression, with the emergence of a lumino-basal cell phenotype that could define the biological properties of this subtype.

### Tumors with lower ER expression express a distinct basal signature

We sought to explore a larger set of ER-positive tumors, specifically to assess whether lower ER expression was associated with expression of a basal gene signature. We first analyzed 564 ER-positive breast cancer tumor cases from The Cancer Genome Atlas (TCGA), in which their ER*α* expression was quantified using Reverse Phase Protein Array (RPPA), which included both ER+/PR+, and ER+/PR- cases (Fig S3A). These cases were stratified into two groups; ER*α* low (141 cases, bottom quartile of ER*α* expression), and ER*α* high (423 cases, top 75% of ER*α* expression) (Fig 3A) and analyzed their basal gene signature^28^. Unsupervised clustering of all basal signature genes revealed modules with higher relative expression in ER*α* low cases compared to ER*α* high cases (Fig S3B). Supervised clustering of these cases based on ER*α* expression revealed a statistically significant (Fig 3B) upregulation of basal signature gene expression in the ER*α* low cluster (Fig 3C) irrespective of PR status (Fig S3C). Similar results were observed when cases were stratified using *ESR1* mRNA levels instead of ER*α* levels (Fig 3D-F, S3D-F), whereby tumors expressing lower *ESR1* demonstrated an increased expression of genes conferring basal identity, suggesting that tumors gain basal-like traits upon concomitant reduction of ER levels.

**Figure 3:**
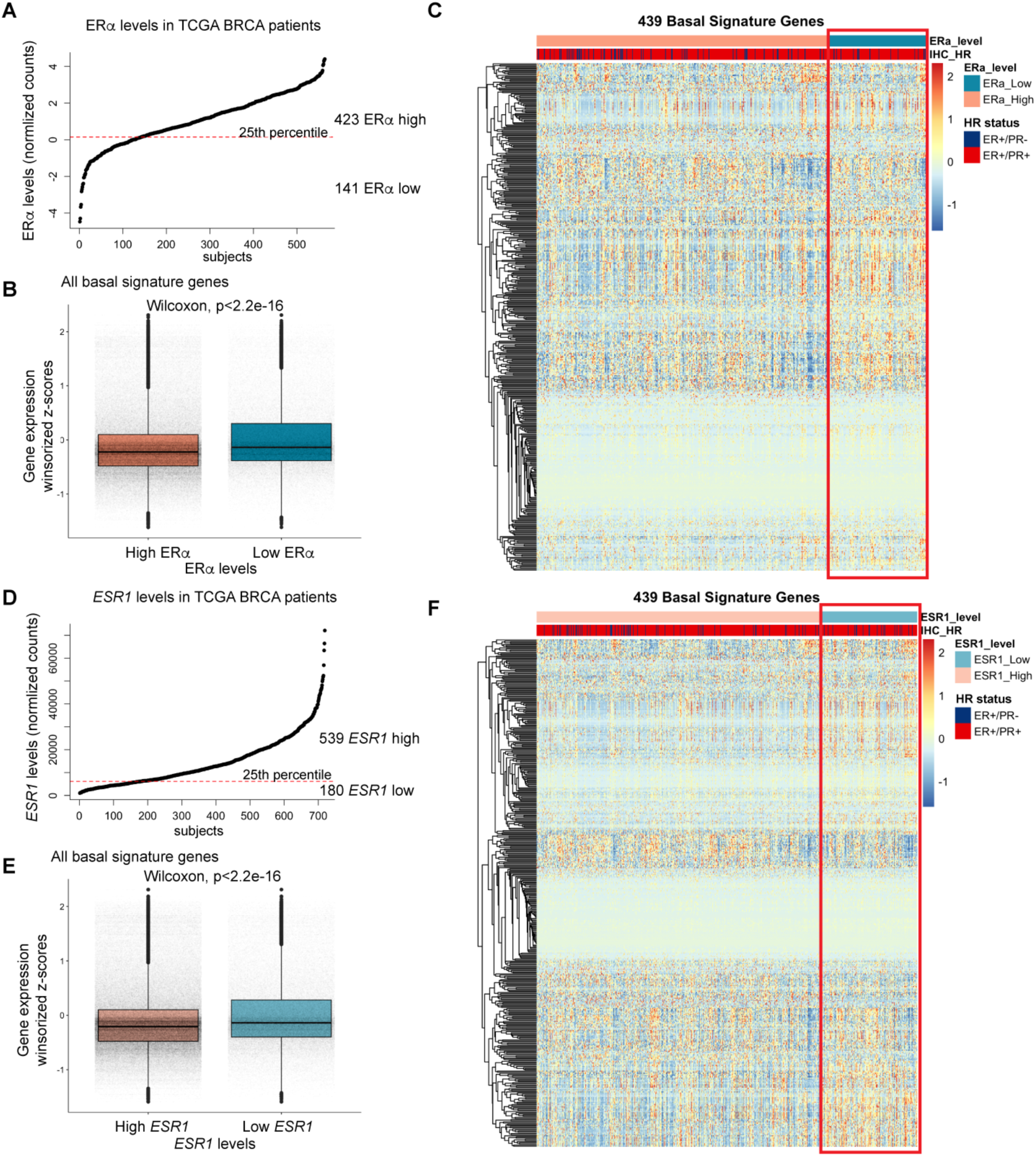
Distinct basal gene expression signature in tumors with lower ER. **(A)** Stratification of ER positive breast tumor cases from TCGA into ER*α* low and ER*α* high groups, based on ER*α* expression. **(B)** Boxplots comparing the distribution of basal signature gene expression in the ER*α* low and ER*α* high groups. **(C)** Heatmap with supervised clustering of the ER positive tumors highlighting higher basal signature gene expression in the ER*α* low group. **(D)** Stratification of ER positive breast tumor cases from TCGA into *ESR1* low and *ESR1* high groups, based on *ESR1* expression. **(E)** Boxplots comparing distribution of basal signature gene expression in the *ESR1* low and *ESR1* high groups. **(F)** Heatmap with supervised clustering of the ER positive tumors highlighting higher basal signature gene expression in the *ESR1* low group.

### Tumor cell plasticity results in emergence of basal-like features in low-ER tumors

We reasoned that the emergence of basal-like characteristics in the low-ER luminal tumors may arise via two possible mechanisms. Firstly, an expansion of basal cells might occur during the later stages of tumorigenesis in low-ER tumors, or alternatively, cellular plasticity may reprogram luminal tumor cells and allow them to acquire basal-like traits.

To identify which mechanism was at play, we carried out lineage tracing using a model that accurately captured aspects of low-ER breast tumors. MMTV-PyMT^29^ is a mouse mammary tumor model that closely resembles human luminal B breast cancers^30^ whereby late-stage tumors lose ER expression^31^. IHC staining for ER revealed weak to moderate expression in MMTV-PyMT tumors, with half of the tumors displaying less that 10% ER expression (Fig S4A and S4B).

In order to trace the lineage of the MMTV-PyMT tumor cells, either Krt8 (luminal) or Krt5 (basal) specific, tamoxifen inducible Cre-ERT promoters^12^ were used to induce expression of GFP in an mTmG reporter mouse^32^ to label luminal or basal cells, respectively (Fig 4A and 4B). The most efficient mammary epithelial GFP labelling was observed when tamoxifen induction was performed 3 days per week on postnatal week 3-old pups for the Krt5-CreERT/Rosa26-mTmG model (Fig S4C), and pups at postnatal weeks 5 and 6 of age for the Krt8-CreERT/Rosa26-mTmG model (Fig S4D). Tumors that eventually arose from these tamoxifen-pulsed mice were harvested and analyzed for GFP expression by immunostaining and flow cytometry.

**Figure 4:**
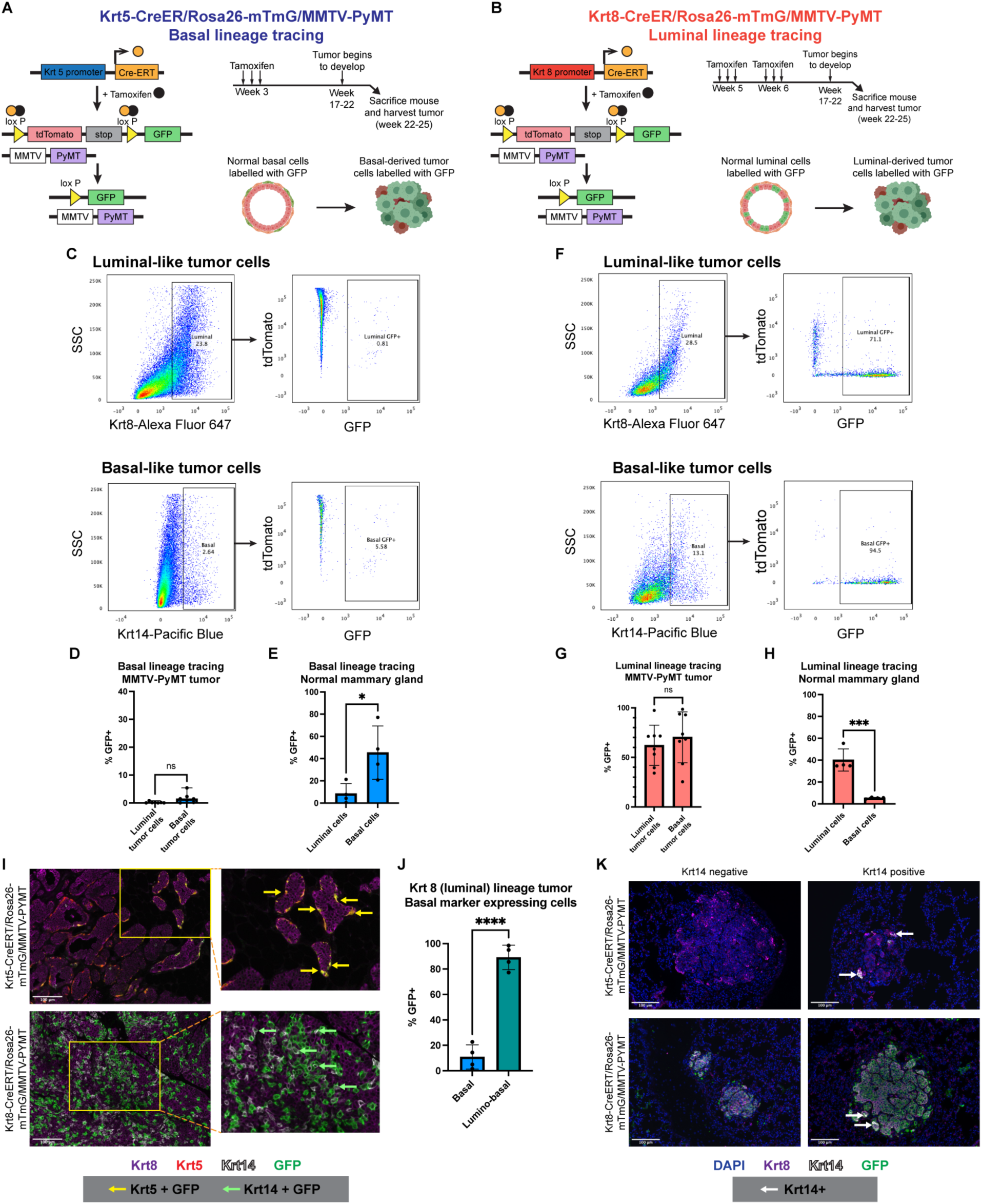
Basal-like MMTV-PYMT tumor cells arise from the luminal lineage. (A-B) Strategy to trace basal and luminal lineage tumor cells in MMTV-PyMT mice. Lineage-specific and tamoxifen-inducible Cre-ERT2 was used to specifically label basal **(A)** or luminal **(B)** epithelial cells with GFP. **(C)** Representative flow cytometry plots of luminal and basal tumor cells inheriting the basal lineage GFP label. **(D-E)** Percentage of basal and luminal cells expressing basal lineage GFP in Krt5-CreERT/Rosa26-mTmG/MMTV- PyMT tumors (n=7, p=0.0513, unpaired t test) **(D)** and the normal Krt5-CreERT/Rosa26- mTmG mouse mammary gland (n=4, p=0.028, unpaired t test) **(E)**. **(F)** Representative flow cytometry plots showing luminal and basal tumor cells inheriting the luminal lineage GFP label. **(G-H)** Percentage of basal and luminal cells expressing luminal lineage GFP in Krt8-CreERT/Rosa26-mTmG/MMTV-PyMT tumors (n=8, p=0.4938, unpaired t test) **(G)** and the normal Krt8-CreERT/Rosa26-mTmG mouse mammary gland (n=4, p=0.0005, unpaired t test) **(H)**. **(I)** Representative images of TSA staining of Krt5-CreERT/Rosa26- mTmG/MMTV-PyMT and Krt8-CreERT/Rosa26-mTmG/MMTV-PyMT tumors. Samples were stained with Krt8 (purple), Krt5 (red), Krt14 (white), and GFP (green). Yellow arrows point to Krt5+/GFP+ tumor cells, and green arrows point to Krt14+/GFP+ tumor cells. **(J)** Quantification of lumino-basal tumor cells expressing GFP and strictly basal tumor cells expressing GFP from TSA-stained images of Krt8-CreERT/Rosa26-mTmG/MMTV-PyMT tumors. **(K)** Representative images of TSA staining of lung metastases from Krt5- CreERT/Rosa26-mTmG/MMTV-PyMT and Krt8-CreERT/Rosa26-mTmG/MMTV-PyMT mice. Samples were stained with Krt8 (purple), Krt14 (white), GFP (green), and DAPI (blue). White arrows point to Krt14-expressing cells.

Flow cytometry analysis (Fig S4E) of tamoxifen pulsed Krt5-CreERT/Rosa26- mTmG/MMTV-PyMT tumors revealed they were comprised of mostly GFP-negative cells (Fig 4C), indicating that MMTV-PyMT tumors did not originate from the Krt5-expressing basal lineage. In addition, most of the basal-marker expressing population (Krt5+ or Krt14+) did not inherit the GFP label from the basal lineage (mean GFP-positive basal cells= 1.67%) (Fig 4C and 4D). In contrast, a larger proportion (mean GFP positive basal cells= 45.4%) of basal cells inherited the GFP label in the developing normal mammary gland (Fig 4E). These results indicate that the basal-like tumor cells were not derived from the basal lineage, and that the basal-like traits emerging within MMTV-PyMT tumors did not arise from expansion of normal basal cells during the process of tumorigenesis.

On the other hand, tamoxifen pulsed Krt8-CreERT/Rosa26-mTmG/MMTV-PyMT mice developed tumors that were primarily comprised of GFP-positive cells (Fig 4F), indicating that they originated from the luminal lineage. Strikingly, most of the basal-marker expressing tumor cells were found to have also inherited the GFP label (mean GFP- positive basal cells= 70.26%) (Fig 4F and 4G). In contrast, the GFP expression within the normal mammary gland was confined to a small percentage of basal cells (mean GFP- positive basal cells= 5.37%) (Fig 4H). This indicates that the basal-like cells within these tumors arose from the luminal lineage, providing evidence for luminal-to-basal plasticity whereby luminal cells acquire basal-like traits.

TSA staining was also used to visualize and confirm the expression of GFP in luminal- or basal-like tumor cells. Luminal-like tumor cells were identified by Krt8 expression whereas basal-like tumor cells were identified by either Krt5 or Krt14 expression. In Krt5- CreERT/Rosa26-mTmG /MMTV-PyMT tumors, co-expression of Krt5 and GFP was restricted to cells in the tumor periphery (Fig 4I), suggesting that cells of the basal lineage were confined to the adjacent normal regions of the tumor. Furthermore, Krt5 itself was primarily expressed in these adjacent normal regions, with very few Krt5-expressing cells within the main tumor. On the other hand, Krt14 expression was more abundant throughout the tumor (Fig 4I), indicating that Krt14 could serve as a more appropriate marker to track basal identity within these tumors.

GFP expression was more abundant throughout the Krt8-CreERT/Rosa26- mTmG/MMTV-PyMT tumors (Fig 4I), reflecting their luminal origin, with most of these GFP-expressing cells co-expressing Krt8. Co-expression of GFP and Krt14 was also observed, with about 50% of basal marker expressing cells co-expressing GFP (mean= 47.79%) (Fig S4F), confirming the luminal lineage of these basal-like tumor cells. Interestingly, most of the cells co-expressing GFP and Krt14 also co-expressed Krt8 (Fig 4I), suggesting that these tumor cells do not completely lose their luminal identity, but instead gain a lumino-basal phenotype. Quantification of these phenotypes within the tumor show that about 90% of the basal marker and GFP co-expressing cells were of the lumino-basal phenotype (mean=89.10%), with only about 10% of cells transitioning to a fully basal phenotype without Krt8 co-expression (mean=10.90%) (Fig 4J). These findings are consistent with the results from the flow cytometry analysis of these tumors which show that most of the tumor cells expressing the basal marker Krt5 or Krt14 descended from the luminal lineage.

### Distant metastases are seeded by tumor cells of luminal origin

In addition to influencing the course of therapy, lineage plasticity could also play an important role in promoting metastasis within luminal-like tumors. To investigate whether one lineage is important for metastasis than the other, we analyzed lungs from Krt5- CreERT/Rosa26-mTmG/MMTV-PyMT and Krt8-CreERT/Rosa26-mTmG/MMTV-PyMT tumor bearing mice. TSA staining of these lungs for Krt8, Krt14, and GFP revealed no GFP expression in all 7 metastases from Krt5-CreERT/Rosa26-mTmG /MMTV-PyMT mice (Fig 4K and S4G). In contrast, 16 out of 17 lung metastases from Krt8- CreERT/Rosa26-mTmG /MMTV-PyMT mice express GFP, indicating that the metastatic colony is seeded from a tumor cell of a luminal lineage.

All metastases, regardless of the model they arose from, exhibited Krt8 positivity (Fig S4G), consistent with the luminal nature of the primary tumor. In contrast, Krt14 was only observed in a fraction of the metastases (6 out of 17), primarily on the periphery of the metastatic colony. While this may suggest that luminal-to-basal plasticity may not be important in metastatic seeding, it is plausible that lumino-basal tumor cells may have lost their basal marker expression following metastatic colony formation.

Next, we addressed the requirement for lumino-basal cells for metastatic outgrowth, in contrast to its role in metastatic seeding. In our previous experiments above, lungs were obtained from euthanized mice when they reached tumor burden (∼1.0mm^3^). Tumor burden was typically achieved about 4 weeks after an initial palpable tumor was observed, which may not allow sufficient time for the outgrowth of seeded metastases. In order to allow the lung metastases to continue to grow beyond this point, we surgically resected the primary tumor from a Krt8-CreERT/Rosa26-mTmG /MMTV-PyMT mouse, and allowed the lung metastases to develop until the surgically resected tumor began to regrow. This took an additional 3 weeks, which allowed the lung metastases to develop over a total of 7 weeks. Larger metastases were observed in this case, along with an increase in smaller metastatic nodules. Consistent with previous observations, all the metastases expressed GFP and Krt8, and only 20 out of 34 metastases exhibited Krt14 positivity (Fig S4G). Again, there is a possibility that lumino-basal tumor cells have lost their basal marker expression upon reaching the lung. As we are not able to study the metastatic process at multiple timepoints, it is unknown whether lumino-basal cells are important in the metastatic process.

### Single cell RNA sequencing reveals SOX10 as a key driver of luminal-to-basal plasticity

To identify genetic drivers enabling the emergence of lineage plasticity, we carried out single-cell RNA sequencing to compare the gene-expression profiles of lineage-restricted luminal tumor cells and luminal-derived basal tumor cells. Tumors were harvested from two Krt8-CreERT/Rosa26-mTmG /MMTV-PyMT mice and sorted by flow cytometry (Fig S5A) for all GFP-expressing cells to specifically analyze both luminal- and basal-like tumor cells of luminal origin. Dimensionality reduction with uniform manifold approximation and projection (UMAP) revealed the presence of a large, connected cluster of cells, in addition to a smaller cluster that showed clear separation from the larger cluster. Unsupervised clustering identified a total of 18 clusters (Fig 5A), with similar results observed in the two mouse replicates (Fig S5B). Using previously described gene signatures^33^, individual cells were scored to quantify their activity of luminal-progenitor, mature-luminal, and basal gene expression programs. Most cells scored highly for the luminal-progenitor signature (Fig S5C), in agreement with previous single-cell RNA analyses of MMTV-PyMT tumors^34^, however cells from multiple clusters demonstrated concomitant luminal-progenitor and basal signatures (Fig S5D), suggesting that tumor cells do not fully establish a basal identity, and instead express a combination of both luminal progenitor and basal markers. Cluster 6 demonstrated the greatest combined luminal and basal signature scores, suggesting these cells likely express a lumino-basal phenotype (Fig 5B and 5C). Cluster 13 demonstrated high scores specifically for the mature luminal signature, while clusters 15 and 16 demonstrated high scores specifically for the basal signature, suggesting that these clusters contain luminal- and basal-like tumor cells, respectively (Fig 5B, 5C, and S5E).

**Figure 5:**
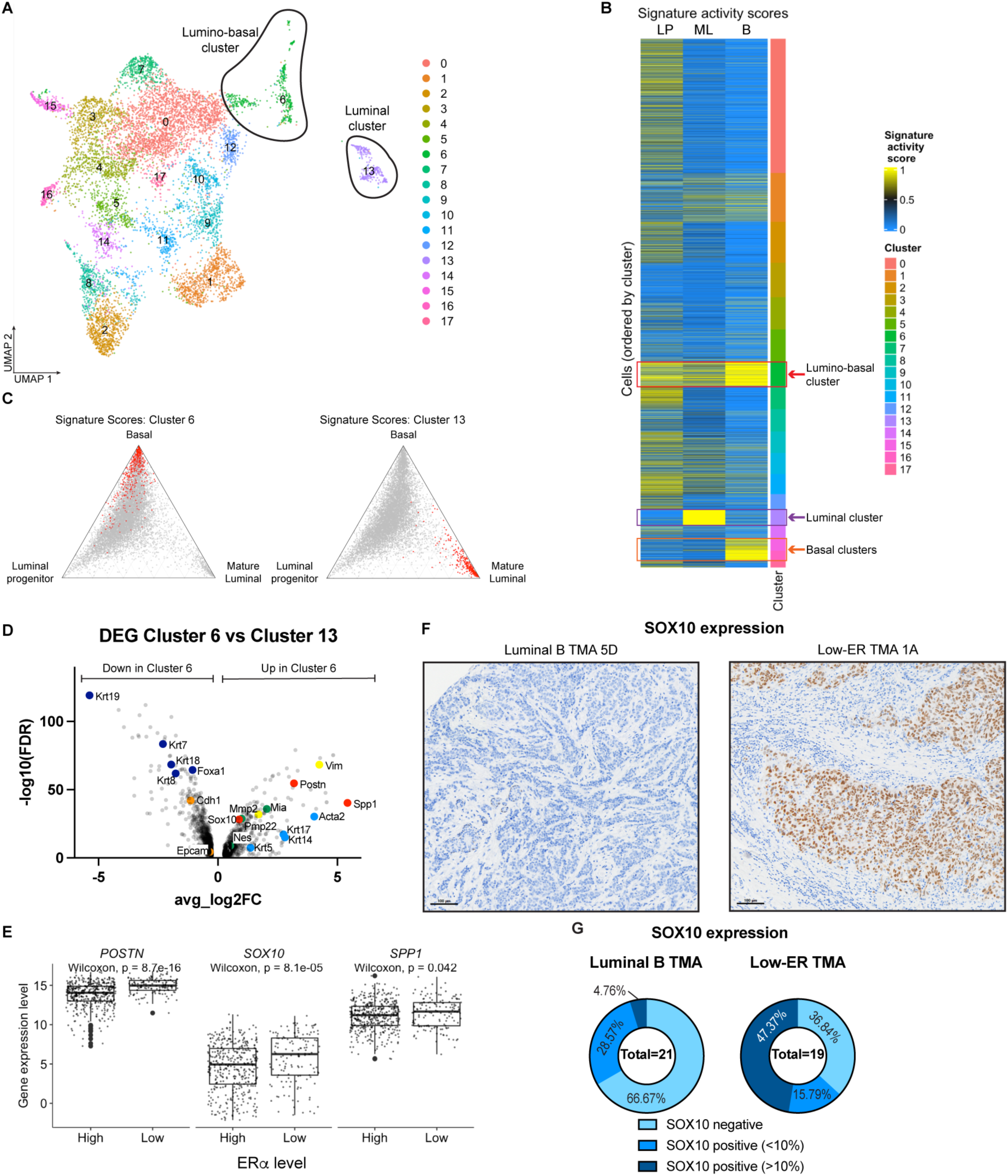
Sox10 is a potential driver of luminal-to-basal plasticity. **(A)** UMAP of Krt8- CreERT/Rosa26-mTmG/MMTV-PyMT tumor cells sorted for GFP. Unsupervised clustering divided the cells into 18 different clusters. **(B)** Heatmap showing the basal, mature luminal, and luminal progenitor gene signature activity in each cluster. Cluster 13 (purple box) has high expression of mature luminal genes, while clusters 15 and 16 (orange box) have high expression of basal genes. Cluster 6 (red box) has high expression of both mature luminal and basal genes, and is identified as the lumino-basal cluster. **(C)** Ternary plots showing the distribution of relative activity of luminal progenitor, mature luminal, and basal gene signatures across all tumor cells profiled by scRNA-seq. In the left plot, cells assigned to cluster 6 are highlighted in red, while cells assigned to cluster 13 are highlighted in the right plot. **(D)** Volcano plot showing genes upregulated and downregulated in cluster 6 as compared to cluster 13. Light blue points are basal marker genes, dark blue points are luminal marker genes, yellow points are mesenchymal genes, orange points are epithelial genes, red points are potential plasticity driver genes, and green points are SOX10 target genes. **(E)** Boxplots showing the differences in distribution of *POSTN*, *SOX10*, and *SPP1* gene expression between the ER*α* low and ER*α* high groups of tumors obtained from TCGA. **(F)** Representative images of SOX10 IHC staining in luminal B and low-ER TMAs. **(G)** Quantified SOX10 expression in luminal B and low-ER TMAs.

To identify genes contributing to this lumino-basal plasticity, we compared the genes expressed in the lumino-basal cell cluster (cluster 6) against their possible ancestors, the lineage-restricted luminal cell (cluster 13). Analysis of the differentially expressed genes between the cluster 6 and cluster 13 uncovered upregulation of basal marker genes (*Krt5*, *Krt14*, *Krt17*, and *Acta2*) (Fig 5D, light blue points) and simultaneous downregulation of luminal marker genes (*Krt7*, *Krt8*, *Krt18*, *Krt19*, and *Foxa1*) (Fig 5D, dark blue points) in cluster 6, supporting the proposed lumino-basal identity of this cluster. Of note, mesenchymal genes *Mmp2*, and vimentin (*Vim*) (Fig 5D, yellow points) were upregulated in cluster 6, while epithelial genes E-cadherin (*Cdh1*) and Epcam (Fig 5D, orange points) were downregulated, suggesting that the emergence of basal-like characteristics in these luminal-derived tumor cells may result from an epithelial to mesenchymal transition (EMT).

From the list of genes upregulated genes in cluster 6, we selected 3 genes that may potentially regulate the luminal-to-basal plasticity observed in low-ER tumors (Fig 5D, red points). The gene *Spp1*, which encodes the protein osteopontin (*Opn*), was selected as it has the highest effect-size (avg_log2FC=5.42). Osteopontin is usually found as a component of bone, as it is an extracellular structural protein, however, intracellular osteopontin has been found to regulate EMT^35^ via its interaction with the stemness marker *Cd44*^36^. We also selected the gene *Postn*, which encodes periostin, an extracellular matrix protein that has been found to enable cell motility by binding to integrins *α*V*β*3 and *α*V*β*5^37^. Finally, we selected the transcription factor *Sox10*, as it has been implicated in the regulation of cell plasticity in mammary tumors^38^. Furthermore, three *Sox10* targets, *Nes*, *Mia* and *Pmp22*, identified using TRRUST v2 (Transcriptional Regulatory Relationships Unraveled by Sentence-based Text mining)^39^, were found to be upregulated in cluster 6 as well (Fig 5D, green points), further supporting the role that this transcription factor may play in driving plasticity.

To assess the roles of each candidate gene on lumino-basal plasticity in low-ER tumors, we analyzed the expression of these genes in the ER*α* low vs ER*α* high stratified tumors obtained from TCGA (Fig 3). The expression of all 3 genes were found to be significantly higher in the ER*α* low group of tumors, as compared to the ER*α* high group (Fig 5E). We further analyzed protein expression of these potential plasticity drivers in the luminal B vs low-ER tumor TMAs by using IHC staining. Periostin expression was mostly relegated to the stroma, with no tumor cells found expressing this protein (Fig S5F). Osteopontin could also be detected in the stroma, while tumor cells staining positive for this protein appear to be present on the edges of the tumor (Fig S5G), however, there was no difference in osteopontin expression between tumors in the luminal B or the low-ER TMAs (Fig S5H). On the other hand, SOX10 appears to have a more direct correlation to low-ER tumors (Fig 5F), with high (>10% of tumor cells) expression of SOX10 observed in 47.47% of low-ER tumors, while only 1 of the luminal B tumors (4.76%) expressed high levels of SOX10. 66.67% of luminal B tumors stained negatively for SOX10, compared to only 38.84% of low-ER tumors (Fig 5G), further indicating that SOX10 is more highly expressed in low-ER tumors, as compared to luminal B tumors. This suggests that SOX10 could be a potential driver of luminal-to-basal plasticity, expressed in both the MMTV-PyMT mouse tumors and in low-ER patient tumors.

## Discussion

Our study demonstrates that low-ER breast carcinomas represent a distinct subset of luminal-like tumors, and should be classified as a separate tumor subtype from luminal A and B tumors for the purposes of therapy. We found that cells within the low-ER tumors undergo luminal-to-basal plasticity, which introduces lumino-basal heterogeneity, allowing them to gain basal-like characteristics. These findings potentially reframe the current intrinsic subtypes of breast cancer as connected to each other, instead of being distinct subsets, and that the different subtypes could essentially represent different stages of evolutionary progression of breast tumors as they deviate from their lineage-of-origin. This lineage divergence may influence breast cancer diagnosis and treatment, as luminal-like tumors which typically exhibit better response to therapy, may evolve into more basal-like counterparts in response to drug-induced evolutionary pressures that could selectively eliminate their luminal-like ancestors while sparing basal-like tumor cells.

Lineage plasticity has also been previously described in other breast carcinoma studies. Li et al^40^ found basal marker expression in luminal lineage-labelled cells, although they did not carry out further investigations on this observation. Hein et al^41^ observed that oncogenic transformation of the luminal lineage resulted in a small percentage of tumor cells co-expressing Krt5, however, their model utilized HA-tagged Polyoma Middle T antigen (PyMT) and ErbB2 oncogenes to both induce transformation and label the luminal lineage, whereas our lineage-tracing strategy uncoupled the process of oncogenesis from lineage labelling. The use of the MMTV promoter to drive the expression of the PyMT oncogene allowed this protein to be expressed in both luminal and basal cells, as previously shown^34^, while lineage labelling using keratin-driven *Cre* recombination, allows specific labelling of either luminal or basal cells. Consistent with our observations, they also found Krt5 expression to be restricted to the edges of the tumor. Finally, Koren et al^42^ and Van Keymeulen et al^43^ noted that PIK3CA mutations could induce oncogenesis and multipotency within mammary cells, resulting in heterogeneous, multi-lineage tumors. These findings suggest that oncogenesis and lineage plasticity can occur simultaneously by the activity of various oncogenes. While we have identified SOX10 as the potential driver for lineage plasticity, it is unknown whether this transcription factor itself is activated by PyMT the driver oncogene used in our model. PyMT is known to induce oncogenic transform via interacting with, and activating c-Src^44^, a non-receptor tyrosine kinase, thereby activating various other signaling molecules such as Shc^45^ and PI3K^46^. It is thus possible that one of the cell signaling pathways activated may lead to an increase in SOX10 transcription and activity, and the PyMT oncogene may indirectly have an effect on SOX10 function. Further studies must be carried out to in order to identify if and how lineage plasticity can occur independently from oncogenesis, as the homogeneous tumor cell populations observed within some luminal-like tumors suggest that oncogenic transformation may not always lead to lumino-basal plasticity. Furthermore, our study showed specific unidirectional luminal-to-basal plasticity, and not simply multipotency, which has important implications when considering the potential evolution from a luminal-like to a basal-like tumor, as mentioned previously.

The presence of basal-like cells within breast tumors has important implications in tumor development and metastasis. We have previously shown that PKA-induced reduction of the lumino-basal subpopulation may be important in limiting metastasis and reducing chemotherapy resistance^34^. The collective migration of breast cancer has also been shown to involve leader cells with a re-activated basal program^47^, suggesting that the successful establishment of metastasis by these invading tumor clusters may depend on leader cells that have undergone luminal-to-basal plasticity. It is important to note that, while we are not able to assess the requirement for lumino-basal plasticity in metastasis from our data, we cannot rule out the possibility that the cells that established these metastases could have gained lumino-basal features within the primary tumor, which may have been subsequently lost upon lung colonization. Metastatic dissemination has been shown to involve cells that have undergone partial EMT^48^, suggesting that maintaining cellular plasticity is beneficial in metastasis. Circulating tumor clusters have also been found to consist of multiple clones^49^, suggesting that only a small percentage of lumino-basal cells may be required to successfully colonize distant sites. Further studies analyzing circulating tumor clusters in lineage-labelled mice, or studying early vs late metastases may help to address the importance of lumino-basal plasticity in metastasis. Alternatively, lineage ablation experiments can be carried out using mouse models with inducible and lineage-specific diptheria toxin, to eliminate the lumino-basal population, which could assess if metastases can still develop without lumino-basal tumor cells.

Although the MMTV-PyMT mouse is an appropriate mammary tumor model which is commonly used in the study of breast carcinomas^31^, the oncogenic mechanism and development of these murine mammary tumors may not fully reflect the typical progression of human carcinomas. The use of this model in our lineage tracing experiments may thus be a possible limitation in attempting to elucidate the origin of lumino-basal heterogeneity within low-ER tumors. The model, however, represents the closest to modeling low-ER breast cancers and to understand lineage plasticity and lumino-basal heterogeneity. Besides the use of this specific mouse model, alternative models may also be useful in attempting to study lumino-basal heterogeneity. Specifically, the *Brca1*^F22–24/F22–24;p53+/−^ mouse tumor model^50^ more accurately reflects mammary epithelial transformation in human patients by introducing *Brca1* mutation and loss of *p53*. When crossed with *BLG-Cre* mice to induce oncogenesis in milk-producing luminal cells, this mouse model was shown to produce tumors that are basal-like and metaplastic. Substitution of *BLG-Cre* with Krt8- or Krt5-driven Cre could enable oncogenic transformation and lineage labelling to occur simultaneously. Lineage labelling of luminal progenitor cells in mice using ELF5-rtta in combination with TetO-Cre^51^ could also help to address whether lumino-basal plasticity can specifically be observed in this luminal subpopulation. Finally, in order to confirm if lumino-basal plasticity is a crucial step in tumor development, lineage ablation experiments can also be carried out to assess whether eliminating the lumino-basal subpopulation would interrupt tumor development.

We have uncovered that SOX10 may be responsible for driving the lumino-basal plasticity seen in the low-ER tumors. This is in agreement with previous studies demonstrating the role of SOX10 in regulating cell-state plasticity in mammary tumors^38^. Interestingly, several recent studies have shown that SOX10 is preferentially expressed in triple negative and metaplastic breast carcinomas and has emerged as a useful immunohistochemical marker to utilize in breast pathology practice^52, 53^. In addition, SOX10 is associated with developmental plasticity and bipotent progenitor identity in fetal mammary stem cells, suggesting that the activity of this transcription factor reflects the reactivation of the bipotent progenitor program in tumor cells. SOX10 has been shown to be expressed in TNBCs, and is associated with worse clinical outcomes in these patients^54^, highlighting the similarities between this tumor subtype and the low-ER tumors in our study. SOX10 has also been shown to induce dedifferentiation and EMT^38^, highlighting the increase in invasive potential of the basal-like progression of low-ER tumors, potentially leading to the worse prognosis and poor clinical outcomes observed in these patients.

## Materials and methods

### Dartmouth-Hitchcock Medical Center pathology database search

The pathology database (Cerner Millennium) at Dartmouth-Hitchcock Medical Center was retrospectively searched from January 2012 through August 2020 to identify all invasive breast cancer cases with low ER expression. Low ER expression was defined as a sample displaying 1-10% of cancer cells with ER expression by immunohistochemistry (IHC), according to the American Society of Clinical Oncology (ASCO) – College of American Pathologists (CAP) 2020 guidelines^8^. Pathology reports were reviewed to include all primary invasive breast cancers with low ER expression, and H&E and IHC slides were reviewed by a breast pathologist (KM). Pathologic characteristics were recorded from pathology reports and slide review and included tumor histologic type, tumor size, tumor grade (Nottingham combined histologic grade/modified Scarff-Bloom-Richardson grade), presence of ductal carcinoma in-situ (DCIS), lymphovascular invasion, axillary lymph node status, and response to neoadjuvant therapy, when administered. For patients with a pathologic complete response after neoadjuvant therapy, tumor characteristics were assessed on the pre-treatment core biopsy. A tumor was considered to exhibit basal-like histologic features when all of the following were present: solid sheets of tumor cells with a syncytial growth pattern, high-grade, pleomorphic cytological features, abundant mitotic activity, tumor circumscription with pushing borders, prominent intratumoral and/or peripheral lymphocytic infiltrates, and tumor necrosis. Patient clinical features including age at diagnosis, treatment regimens, and follow-up status were recorded from electronic medical records.

### Determination of ER, PR, and *HER2/neu* expression

ER, PR, and *HER2/neu* were performed on diagnostic core needle biopsies in all cases. Biomarkers were repeated on a subsequent surgical specimen at the request of treating clinicians in a minority of cases (n = 10). Immunohistochemical assays for ER and PR were performed on paraffin-embedded tissue sections fixed in 10% neutral buffered formalin for 6-72 hours using the polymer system technique with appropriate controls. The assays were performed according to the manufacturer’s instructions using Anti-ER (Cell Marque, 249R-15-ASR, clone: SP1) and Anti-PR (Biocare Medical, ACA424B, clone: 16) antibodies. ER and PR were qualified (positive or negative) and quantified (% of tumor cells staining) by breast pathologists by “eyeballing” IHC stained slides. In addition, we evaluated for ER staining intensity (weak, moderate, or strong). *HER2/neu* analysis was performed using dual-probe FISH (Abbott Laboratories, PathVysion HER-2 DNA probe kit) to assess for gene amplification and results were interpreted in accordance with the ASCO/CAP HER2 testing guidelines^55^.

### Animal studies

All animal experiment IACUC protocols were approved by the Dartmouth College Committee on Animal Care. MMTV-PyMT mice ((Tg(MMTV-PyVT)^634Mul/LellJ^ mice on a C57Bl/6J background, strain #: 022974)^29^, Krt5-CreER mice (B6N.129S6(Cg)- *Krt5^tm1.1(cre/ERT2)Blh^*/J, strain #: 029155)^12^, Krt8-CreER mice (Tg(Krt8-cre/ERT2)^17Blpn^/J, strain #: 017947)^12^, and Rosa26-mTmG reporter mice (B6.129(Cg)- Gt(ROSA)26Sor^tm4(ACTB-tdTomato,-EGFP)Luo^/J, strain #: 007676)^32^ were purchased from The Jackson Laboratory. For tamoxifen induced mammary epithelial labelling, tamoxifen (Sigma-Aldrich, T5648-1G) was prepared by dissolving in commercially available corn oil for 5 hours at 37°C to a final concentration of 30mg/ml. Krt5-CreERT/Rosa26-mTmG /MMTV-PyMT mice were administered 150mg/kg (100ml of tamoxifen stock for a 20g mouse) of the tamoxifen stock 3 times per week at week 3, while Krt8-CreERT/Rosa26- mTmG /MMTV-PyMT mice were administered 150mg/kg of the tamoxifen stock, 3 times per week at weeks 5 and 6. Mice were euthanized and tumors were harvested once tumors reached a volume of 1.5cm^3^, usually at weeks 20-25. For analysis of the normal mammary gland, mice were euthanized at 8 weeks or age matched to tumor bearing mice.

### Mammary gland dissociation

Mouse mammary fat pads were harvested and processed to obtain single-cell suspensions using established protocols^56^ that were slightly modified. Mammary fat pads were digested in a solution of DMEM (Corning, 10-013-CV) with Hyaluronidase (Fischer Scientific, ICN10074091) and Collagenase A (Sigma-Aldrich, 10103586001) for 2 hours at 37°C with gentle agitation using a rotator. Red blood cells were subsequently removed with an ammonium chloride lyse (8.02g NH4Cl, 0.84g NaHCO3, 0.37g EDTA in 1L of water), and samples were agitated with Trypsin (Corning, 25-053-CI) and Dispase (Stem Cell Technologies, 7913) + DNAse I (Sigma-Aldrich, DN25-100mg) for 1 minute each to further dissociate the cells. Finally, samples were filtered through a 40mm cell strainer (Corning, 431750) to obtain a single-cell suspension.

### Mammary gland whole mount preparation and Carmine Alum staining

Whole mammary glands were spread on a glass slide and fixed with Carnoy’s fixative (60% ethanol, 30% chloroform, 10% glacial acetic acid) overnight at RT. Fixed tissue was rehydrated by washing with decreasing ethanol concentrations (70%, 50%, 30%, 10%) 2 times each for 10 minutes. Rehydrated tissue was then stained with Carmine Alum (Stem Cell Technologies, 07070) for 48-72 hours. Mammary glands were then dehydrated using increasing ethanol concentrations (70%, 95%, 100%) 2 times each for 15 minutes, and cleared in xylene overnight. Cleared mammary glands were then mounted with Permount mounting medium (Fischer Chemical, SP15-100) and glass coverslips and allowed to dry overnight. Slides were imaged on the PerkinElmer Vectra3 slide scanner.

### Tumor dissociation

Tumors harvested from euthanized mice were digested in DMEM containing 2 mg/ml Collagenase A and 100U/ml hyaluronidase at 37°C for 2 hours with gentle agitation using a rotator. Following digestion, samples were strained through 70mm (Corning, 431751) and 40mm cell strainers to obtain a single-cell suspension. Finally, red blood cells were removed with an ammonium chloride lyse, and cells were washed in PBS.

### FFPE tissue processing

Harvested mammary glands, tumors and lungs were placed in tissue biopsy cassettes and fixed in 10% Neutral Buffered Formalin (Leica, 3800598) at 4°C overnight. The formalin was then removed and tissues were soaked in 70% ethanol at 4°C for at least 2 days before embedding in paraffin blocks. Hematoxylin & Eosin (H&E) staining was performed on sections cut from the paraffin blocks. Embedding, sectioning, and H&E staining were performed by Dartmouth-Hitchcock Pathology Shared Resources.

### Flow Cytometry and Fluorescence assisted cell sorting (FACS)

Single-cell suspensions were first stained with fluorescently labelled antibodies. Tumor single-cell suspensions were stained with Alexa Fluor 700 anti-CD326 (Ep-CAM) antibody (Biolegend, 118239, clone: G8.8, 1:100 dilution), PE/Cyanine 7 anti-mouse CD31antibody (Biolegend, 102418, clone:390, 1:100 dilution), and PE/Cyanine 7 anti-mouse CD45 antibody (Biolegend, 103114, clone: 30-F11, 1:100 dilution) for 30 minutes on ice. Mammary gland single-cell suspensions were stained with all the above antibodies, with the addition of Super Bright 600 anti-CD49f (integrin alpha 6) antibody (Thermo Scientific, 63-0495-42, clone: GoH3, 1:100 dilution).

For staining intracellular keratins, single-cell suspensions were fixed for 15 minutes at RT in 2% paraformaldehyde (methanol free, Thermo Scientific, J19943-K2), and permeabilized for 15 minutes at RT in Intracellular Staining Perm Wash Buffer (Biolegend, 421002), before staining with Recombinant anti-Cytokeratin 8 antibody Alexa Fluor 647 (Abcam, ab192468, clone: EP1628Y, 1:100 dilution), and Recombinant anti-Cytokeratin 5 antibody (Abcam, ab236216, clone: SP27) conjugated to Dylight 405 (Abcam, ab201798), or anti-Cytokeratin 14 monoclonal antibody (Thermo Scientific, MA5-11599, clone: LL002), conjugated to Pacific Blue (Thermo Scientific, P30013) for 30 minutes at RT. Samples were washed and resuspended in PBS supplemented with 2% FBS before being analyzed for cell marker expression using BioRad ZE-5 cell analyzer.

Compensation was performed with the aid of single-stained Ultracomp eBeads plus compensation beads (Invitrogen, 01-3333-42). Analysis and plot generation was performed on FlowJo.

For sorting of GFP-expressing cells, single-cell suspensions were only stained for extracellular markers, and DAPI (Sigma-Aldrich, 10236276001) was added at a dilution of 1:1000 after the final wash step in order to facilitate live-cell sorting. GFP-positive cells were sorted on FACSAria III cell sorter by first gating on DAPI-negative live cells, and CD31- and CD45-negative epithelial cells.

### Immunohistochemistry (IHC) staining

Slides are cut at 4mm and air dried at RT before baking at 60°C for 30 minutes. Automated protocol performed on the Leica Bond Rx (Leica Biosystems) includes paraffin dewax, antigen retrieval, peroxide block and staining. Heat induced epitope retrieval using Bond Epitope Retrieval 2, pH9 (Leica Biosystems, AR9640) was incubated at 100 degrees Celsius for 20 minutes (for anti-Cytokeratin 5, Bond Epitope Retrieval 1, pH6.0 (Leica Biosystems, AR9961) was used instead). Primary antibody anti-ER (Cell Marque, 249-R-15-ASR, clone: SP1, 1:35 dilution), anti-p63 (Biocare Medical, CM163B, clone:4A4, 1:200 dilution), and anti-Cytokeratin 5 (Abcam, ab236216, clone: SP27, 1:100 dilution) was applied and incubated for 15 minutes at room temp. Primary antibody binding is detected and visualized using the Leica Bond Polymer Refine Detection Kit (Leica Biosystems DS9800) with DAB chromogen and Hematoxylin counterstain. Slides were imaged using the PerkinElmer Vectra3 slide scanner, and PhenoChart. Staining was qualified and quantified by a breast pathologist (KM).

### Multiplexed TSA staining

Staining was optimized based on the PerkinElmer OPAL Assay Development Guide (August 2017). Sample slides were baked at 60°C for 2 hours to remove paraffin wax, followed by 3 xylene washes of 10 minutes each. Slides were then rehydrated with decreasing concentrations of ethanol (100%, 95%, 70%, and 50%), followed by fixation in 10% Neutral Buffered Formalin (Leica, 3800598) for 30 minutes at RT. Antigen retrieval was performed in BOND Epitope Retrieval Solution 1 (Leica, AR9961) for 20 minutes at high pressure in a pressure cooker. After the slides were cooled, they were rinsed in PBS, and endogenous peroxide activity was blocked by treatment with 3% hydrogen peroxide for 10 minutes. Slides were washed in TBS + 0.1% tween (TBS-T) and blocked in Antibody Diluent/Block (Akoya Biosciences, ARD1001EA) for 30 minutes at RT. Primary antibody were added and slides were incubated at RT for 30 minutes. After TBS-T washes, secondary antibodies were added and incubated at RT for another 30 minutes, followed by TBS-T washes. Opal fluorophore was applied to slides for precisely 6 minutes, followed by TBS-T washes. Slides were then boiled for 2 minutes in a microwave at 100% power, followed by 15 minutes at 20% power in AR6 Buffer (Akoya Biosciences, AR600250ML) to affix Opal to target sites and remove primary and secondary antibodies. This process is repeated for each primary antibody used. After staining with the final antibody, Spectral DAPI (Akoya Biosciences, FP1490) was added, and slides were mounted with ProLong Diamond Antifade Mountant (Invitrogen, P36961) and glass coverslips.

The primary antibody and Opal pairs used are as follows:

For TMA samples: anti-Cytokeratin 14 (Abcam, ab119695, clone:SP53, 1:200 dilution) with Opal 620 (Akoya Biosciences, FP1495001KT, 1:500 dilution), anti-Cytokeratin 5 (Abcam, ab64081, clone: SP27, 1:300 dilution) with Opal 520 (Akoya Biosciences, FP1487001KT, 1:150 dilution), anti-ER (Cell Marque, 249-R-15-ASR, clone: SP1, 1:70 dilution) with Opal 650 (Akoya Biosciences, FP1496001KT, 1:500 dilution), and anti- Cytokeratin 8 (Abcam, ab53280, clone: EP1628Y, 1:400 dilution) with Opal 570 (Akoya Biosciences, FP1488001KT, 1:600 dilution).

For mouse tumor and lung samples: anti-Cytokeratin 14 (1:200 dilution) with Opal 620 (1:500 dilution), anti-Cytokeratin 5 (1:300 dilution) with Opal 690 (Akoya Biosciences, FP1497001KT, 1:150 dilution), anti-GFP (Cell Signaling, 2956, clone: D5.1, 1:150 dilution) with Opal 650 (1:500 dilution), and anti-Cytokeratin 8 (1:300 dilution) with Opal 570 (1:600 dilution).

### Image processing, analysis, and phenotype training

Whole slide scans were imaged at 4x resolution using the PerkinElmer Vectra3 slide scanner, and Regions of interest (ROIs) were selected on PhenoChart. ROIs were then imaged at 20x resolution. Spectral unmixing was performed, and each Opal was assigned a color using the software InForm, which was also used to train the algorithm for phenotype quantification. Tissue and cell segmentation was performed (with the aid of DAPI as the nuclear marker, and Krt8, as the cytoplasmic marker), and cells were phenotyped based on marker expression, and validated by marker distribution (entire Cell Mean Fluorescent units extracted for each marker and normalized as a percentile of maximum and minimum fluorescence across all cells in all images).

### Re-analysis of TCGA breast cancer cohort

Preprocessed protein expression data from RPPA assays and gene expression data from RNA-seq from breast cancer patients in The Cancer Genome Atlas (TCGA) were downloaded from Synapse (https://doi.org/10.7303/syn300013).

For ER*α* based analyses, 564 subjects of Luminal A and B primary breast tumors that were estrogen receptor-positive (ER+) with defined progesterone receptor (PR) status from immunohistochemistry staining were used. Binary classification of ER*α* levels were determined based on the distribution of ER*α* from RPPA assays. Subjects were defined as ER*α* low (n = 141) if the level of ER*α* were below the 25^th^ percentile and as ER*α* high (n = 423) if the level of ER*α* were above the 25^th^ percentile.

For *ESR1* based analyses, 719 subjects of Luminal A and B primary breast tumors that were ER+ with defined PR status from immunohistochemistry staining were used. Binary classification of *ESR1* levels were determined based on the distribution of *ESR1* from RNA-seq. Subjects were defined as *ESR1* low (n = 180) if the level of *ESR1* were below the 25^th^ percentile and as *ESR1* high (n = 539) if the level of *ESR1* were above the 25^th^ percentile.

Gene expression levels of 439 basal signature genes (obtained from previously published data^28^) were available in the TCGA data. Winsorized Z-scores for each gene were used to compare the differences in expression between ER*α*/*ESR1* high and low subjects. Log2 transformed counts were used to compare the differences for *POSTN, SPP1*, and *SOX10*.

### scRNA-seq sample and library preparation

GFP expressing cells were collected from the FACSAria, resuspended in PBS + 0.05% BSA and brought to the Genomics Shared Resource for processing. Cell suspensions were counted on a Nexcelom K2 automated cell counter and loaded onto a Chromium Single Cell G Chip (10x Genomics Inc.) targeting a capture rate of 10,000 cells per sample. Single cell RNA-seq libraries were prepared using the Chromium Single Cell 3’ v3.1 kit (10x Genomics) following the manufacturer’s protocol. Libraries were quantified by qubit and peak size determined by Fragment Analyzer (Agilent). All libraries were pooled and sequenced on an Illumina NextSeq2000 using Read1 28bp, Read2 90bp to generate an average of 25,000 reads/cell.

### scRNA-seq data analysis

Raw sequencing data were demultiplexed to create individual FASTQ files using Cell Ranger (v.6.0.1) *mkfastq* (10X Genomics)^57^. Cell Ranger *count* was used to map sequence reads to the reference genome (mm10-2020-A) and construct a matrix of raw read counts. R-package Seurat (v.4.0.4)^58^ was used for downstream processing, normalization, and dataset integration. Raw read counts for cell-containing droplets were imported into R v.4.0.3 using Seurat function *Read10X*. Doublets were identified and removed using the simulation-based approach implemented by function *scDblFinder* from R-package scDblFinder^59^, with the doublet rate argument (dbr) set based on the number of recovered cells from each experiment. Cells with ≤ 500 UMIs or ≤ 200 detected features were removed from further analyses. Cells were further filtered to exclude those identified as outliers (using function *isOutlier* from R-package scater v.1.18.6^60^) from the distribution of mitochondrial read counts (percentage reads mapped to mitochondrial genes). Genes with <10 assigned reads across all samples were also removed prior to downstream analysis. Read counts were normalized using *sctransform*^61^ with default settings. Datasets were integrated using the anchor-based integration framework implemented in Seurat, using 3000 integration features and reciprocal principal components analysis (RPCA) for anchor selection. Unsupervised clustering was performed at multiple resolutions using default parameters in Seurat.

Clustree v.0.4.4^62^ was used to identify the optimal clustering solution for downstream analysis. Contaminating clusters expressing markers of lymphoid and myeloid lineages were identified and removed from the dataset, and the remaining cells were resubjected to unsupervised clustering analysis and dimensionality reduction. Cluster specific marker genes for each cluster were identified using the Seurat function *FindMarkers* with arguments “min.pct = 0.1, only.pos = TRUE” and default parameters. Differential expression analysis was also performed using *FindMarkers* with argument “min.pct = 0.1” and default parameters. Luminal progenitor (LP), mature luminal (ML), and basal (MS) gene expression signatures (obtained from Pal *et al.*^33^) were scored for enrichment at the individual cell-level using variance-adjusted Mahalanobis (VAM)^63^. VAM generates cell-specific scores, using the gamma cumulative distribution function (CDF), between 0 (no enrichment) and 1 (highly enriched) for a given gene-set. Log-normalized counts (generated by Seurat function *NormalizeData*) were used as input to function *vamForSeurat,* which was run with default settings. Squared adjusted Mahalanobis distances were used to generate ternary plots, positioning each cell according to its combined expression of LP, ML, and MS gene signatures.

### Raw scRNA-seq data

The data discussed in this publication have been deposited in NCBI’s Gene Expression Omnibus (Edgar et al., 2002^64^) and are accessible through GEO Series accession number GSE214815 (https://www.ncbi.nlm.nih.gov/geo/query/acc.cgi?acc=GSE214815).

## Acknowledgements

Part of the illustrations in Figure 4 were created with BioRender.com. We thank Gary Ward, Jennifer Fields, Scott M. Palisoul, Dr Meredith Brown, Dr Pat Robison and Dr Fred W. Kolling IV, for sharing their technical expertise, and Dartmouth Center for Comparative Medicine and Research (CCMR) for animal husbandry services. We acknowledge the following Shared Resources facilities: Dartlab (flow cytometry), Pathology shared resources, Microscopy shared resources, Genomics and Molecular Biology Shared Resource, and Dartmouth Cancer Center shared equipment, at the Dartmouth Cancer Center with the NCI Cancer Center Support Grant 5P30 CA023108-43. Single cell studies were conducted through the Dartmouth Center for Quantitative Biology (COBRE, 5P20GM130454-03), in collaboration with the GMBSR with support from NIGMS (P20GM130454) and NIH S10 (S10OD025235) awards. This study was supported by awards 5R00CA201574-05 and 1R01CA267691 from the NIH to D.R.P.

## Author contributions

This project was conceived and designed by G.A.M and D.R.P. Mouse experiments were carried out by G.A.M and N.B.O. Pathology database search and compilation of human patient data was performed by S.M and K.E.M. Analysis of TCGA data was performed by M.K.L and B.C.C. G.A.M carried out all other experiments. Single-cell RNAseq data was analyzed by O.M.W with input from G.A.M. This manuscript was prepared by G.A.M and D.R.P, with contributions from all co-authors.

## Declarations

### Ethical Approval

Research involving animals was carried out in accordance with The Institutional Animal Care and Use Committee (IACUC) and the Dartmouth College Committee on Animal Care (IACUC approval number: 2119).

### Availability of data and materials

The data discussed in this publication have been deposited in NCBI’s Gene Expression Omnibus and are accessible through GEO Series accession number GSE214815 (https://www.ncbi.nlm.nih.gov/geo/query/acc.cgi?acc=GSE214815).

