## Supplementary Figures for "Lineage plasticity enables low-ER luminal tumors to evolve and gain basal-like traits"

**A**

| Histological features of 46 Low-ER positive carcinomas | |
|---|---|
| Type | Invasive ductal carcinoma: 96% (44)<br>Invasive lobular carcinoma: 4% (2) |
| Grade | High: 89% (41)<br>Intermediate: 11% (5) |
| Size | 2.97 ± 2.6 cm (range: 0.1 – 10.0 cm) |
| Ductal carcinoma in-situ present | 55% |
| Lymphovascular invasion present | 39% |
| Lymph node status | N0 = 30 (ITCs in two patients)<br>N1 = 2 N2 = 0 N3 = 4 |
| Response to neoadjuvant chemotherapy | Pathologic complete response = 75% (9/12)<br>Pathologic partial response = 25% (3/12)*<br>*Two had microinvasion only |

Oncotype Recurrence Score (n=10) = 48.8 ± 12 (18-61)

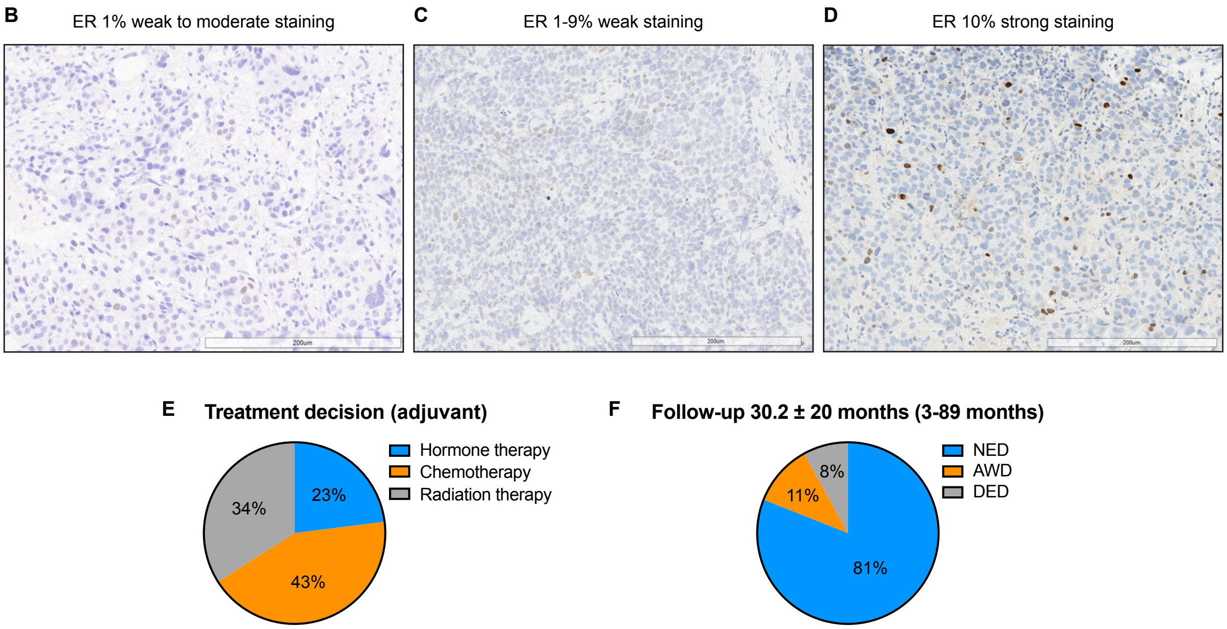


**Supplementary Figure 1: (A)** Histological features of the 46 low-ER cases in our cohort. **(B-D)** Representative IHC images showing the different levels and intensity of ER staining in the low-ER tumors. **(E-F)** Breakdown of treatment decisions **(E)**, and follow-up details **(F)**, of the 46 low-ER cases.

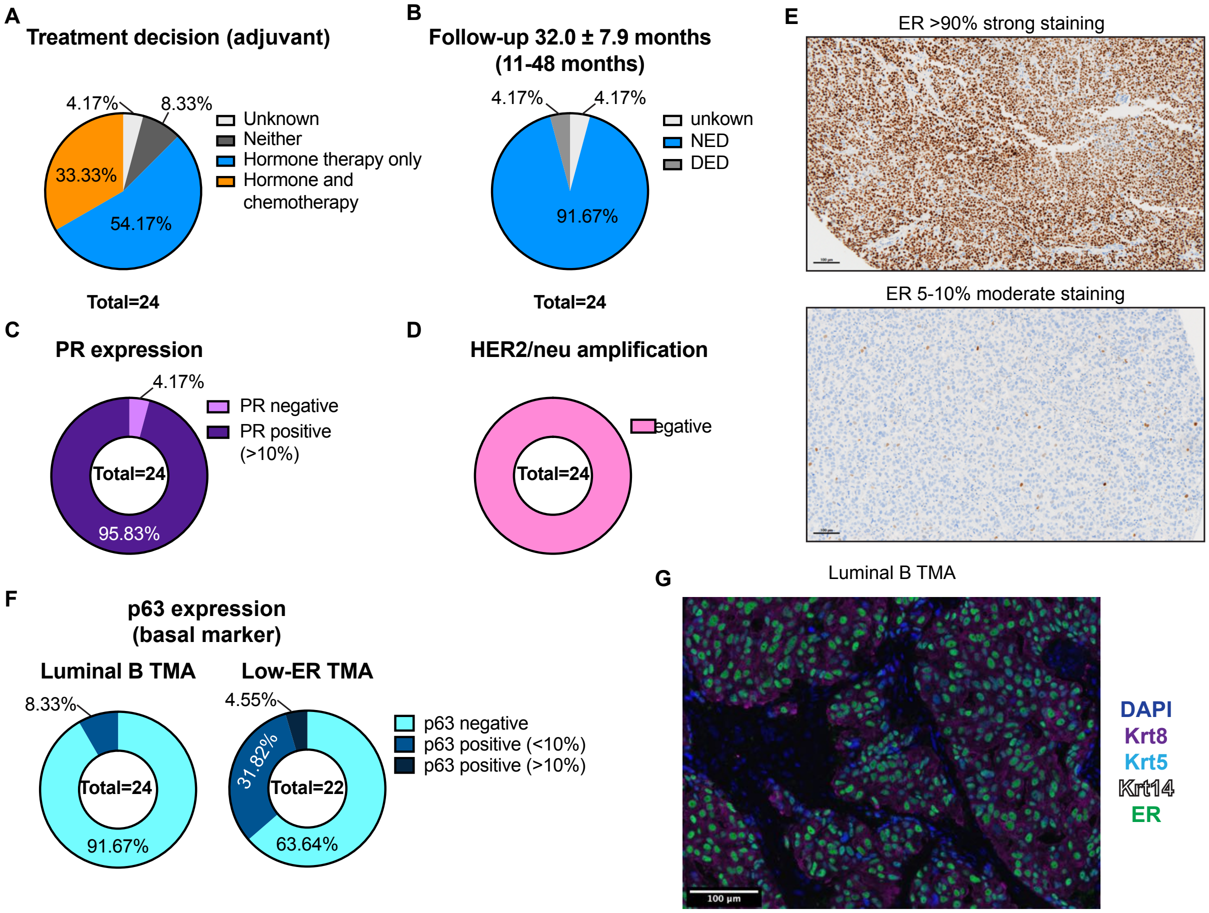


**Supplementary figure 2: (A-D)** Breakdown of treatment decisions **(A)**, follow-up details **(B)**, PR expression **(C)**, and Her2/neu amplification **(D)**, of the 24 luminal B cases used in the TMA. **(E)** Representative IHC images showing the different levels and intensity of ER staining in the luminal B TMA cores. **(F)** p63 expression differences between luminal B and low-ER tumors. **(G)** Representative images of TSA staining showing homogeneous populations in luminal B TMAs. Samples were stained with Krt8 (purple), Krt5 (cyan), Krt14 (white), ER (green), and DAPI (blue).

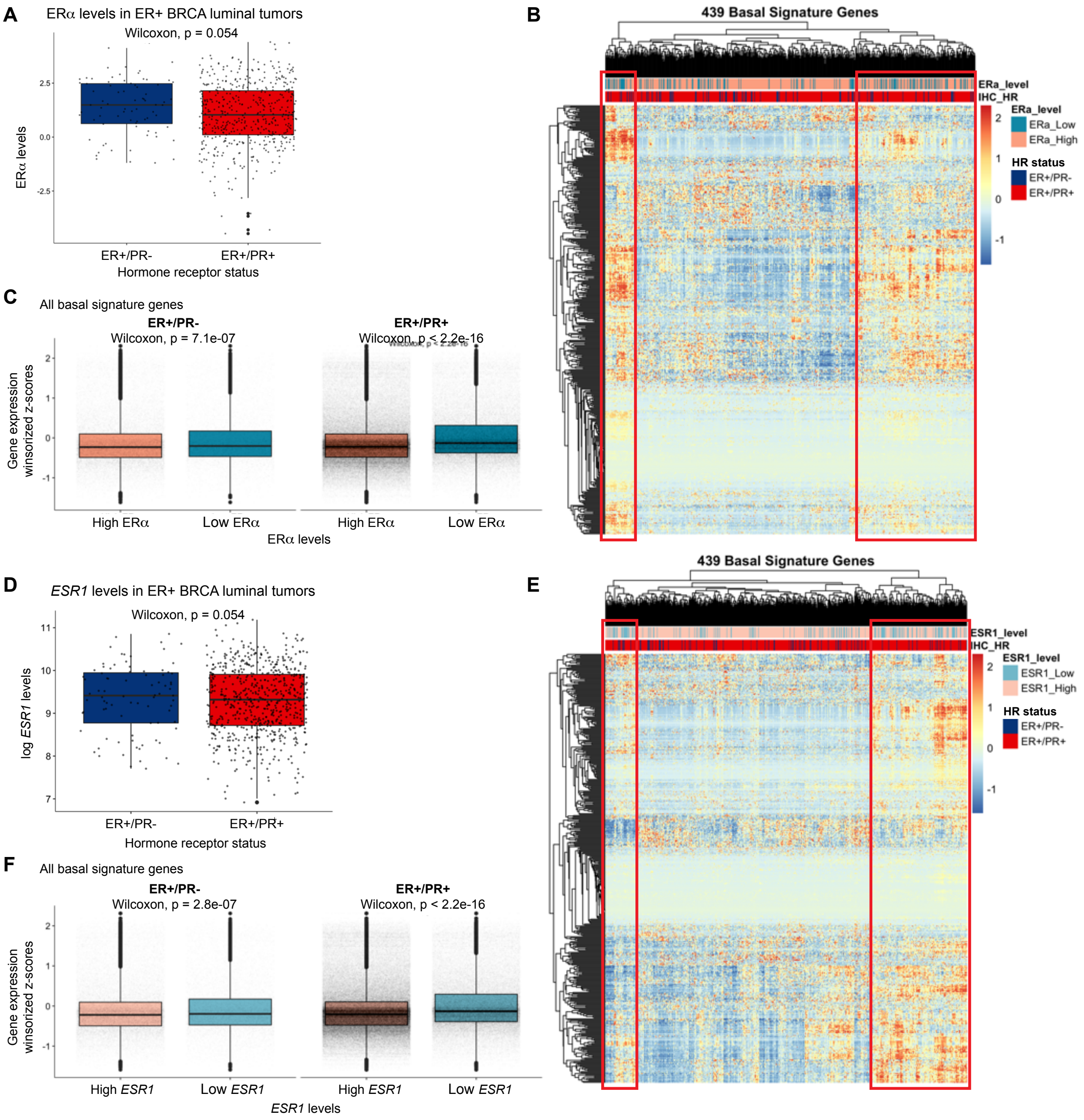


**Supplementary figure 3: (A)** Boxplots showing distribution of ERα levels in the ER+/PR- and ER+/PR+ cases used in this analysis. **(B)** Heatmap with unsupervised clustering of the basal signature gene expression in ER positive tumors. **(C)** Boxplots showing the differences in distribution of basal signature gene expression stratified by ERα expression level and PR status. **(D)** Boxplots showing distribution of *ESR1* levels in the ER+/PR- and ER+/PR+ cases used in this analysis. **(E)** Heatmap with unsupervised clustering of the basal signature gene expression in ER positive tumors. **(F)** Boxplots showing the differences in distribution of basal signature gene expression stratified by *ESR1* expression level and PR status.

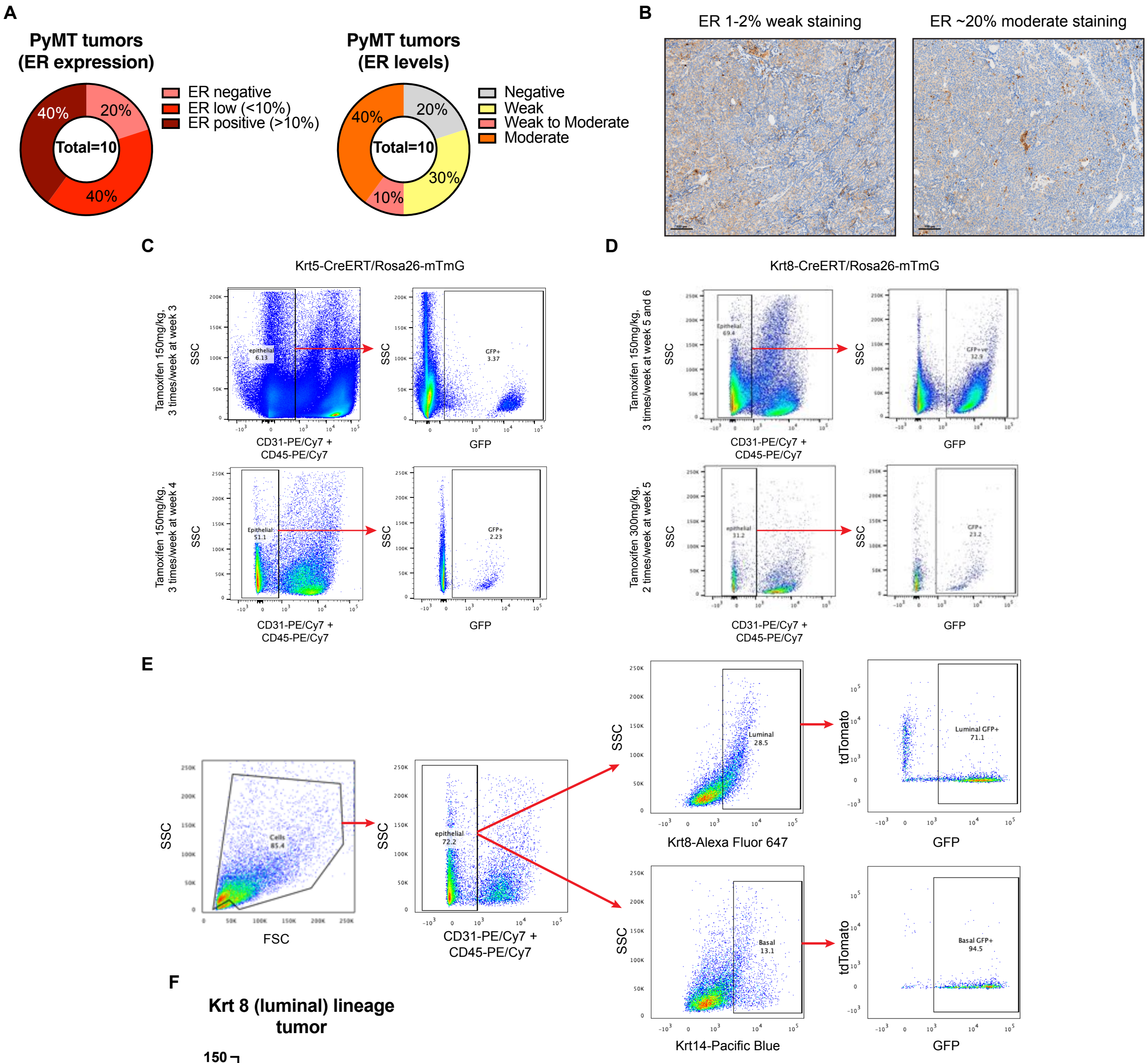


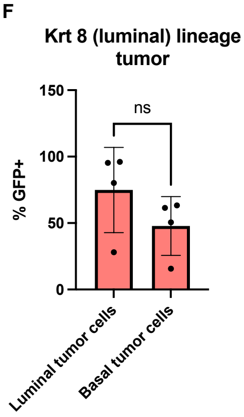


G

| | Krt5-mTmG/ MMTV-PyMT | Krt8-mTmG/ MMTV-PyMT | Krt8-mTmG/MMTV-PyMT (Surgically resected tumor) |
|---|---|---|---|
| Number of mice | 2 | 3 | 1 |
| Total mets | 7 | 17 | 34 |
| Mets expressing GFP | 0/7 | 16/17 | 34/34 |
| Mets expressing Krt8 | 7/7 | 17/17 | 34/34 |
| Mets expressing Krt14 | 4/7 | 6/17 | 20/34 |

**Supplementary figure 4: (A)** ER expression in MMTV-PyMT tumors. **(B)** Representative IHC images of ER expression in MMTV-PyMT tumors. **(C-D)** Optimization of tamoxifen induced GFP labelling in Krt5-CreERT/Rosa26-mTmG **(C)** and Krt8-CreERT/Rosa26-mTmG **(D)** mouse mammary gland. **(E)** Flow cytometry gating strategy for identifying GFP expressing luminal and basal tumor cells. **(F)** Quantification of GFP expressing luminal and basal tumor cells from TSA stained images of Krt8-CreERT/Rosa26-mTmG/MMTV-PyMT tumors. **(G)** Counts of metastatic colonies from Krt5-CreERT/Rosa26-mTmG/MMTV-PyMT and Krt8-CreERT/Rosa26-mTmG/MMTV-PyMT mice expressing Krt8, Krt14, and GFP.

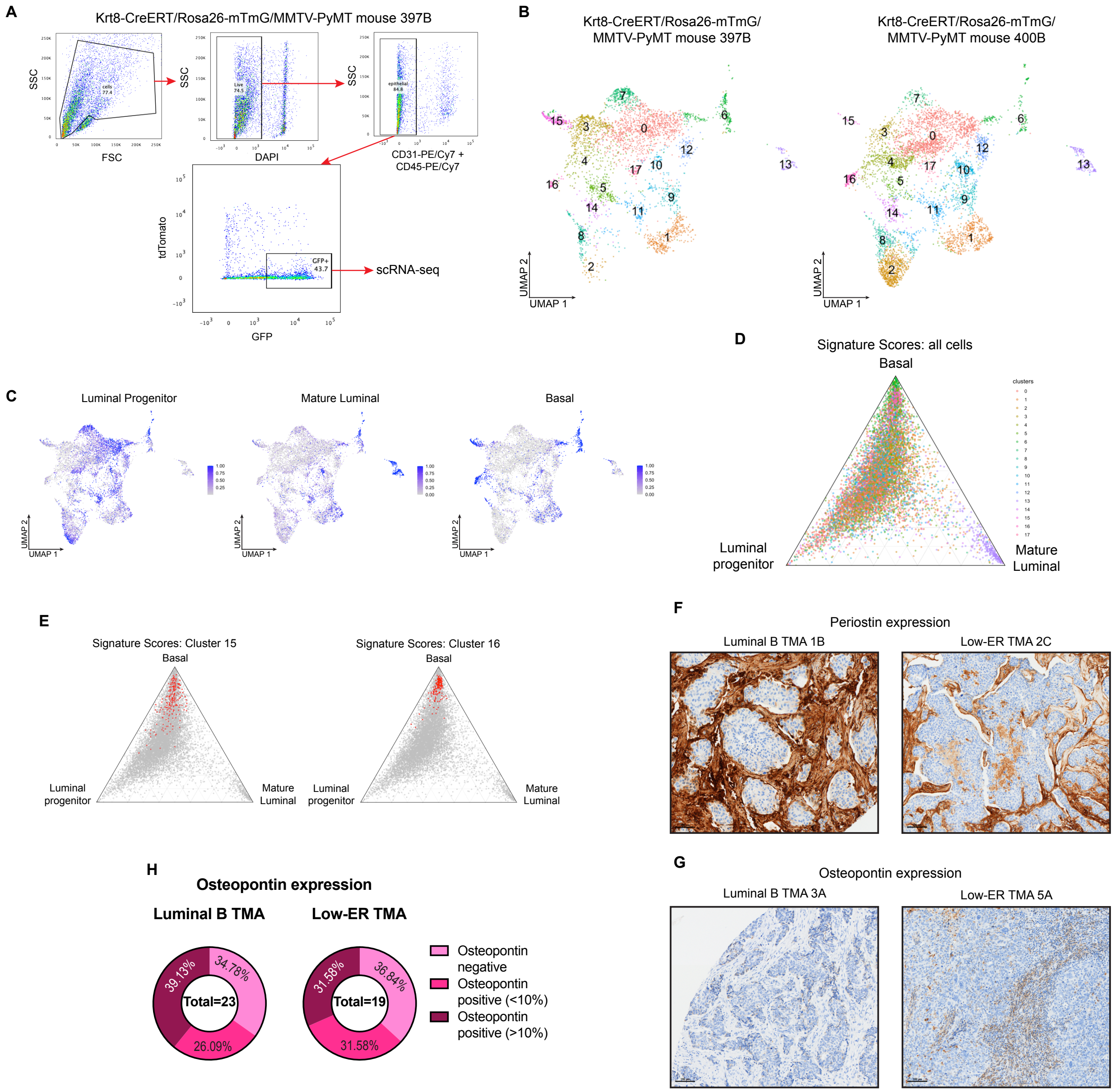


**Supplementary figure 5: (A)** Flow cytometry plots showing the gating strategy and the GFP positive cell population harvested using fluorescence assisted cells sorting (FACS). **(B)** UMAP of Krt8-CreERT/Rosa26-mTmG/MMTV-PyMT tumor cells sorted for GFP and split by sample of origin. Unsupervised clustering divided the cells into 18 different clusters. **(C)** UMAP projections of all tumor cells profiled with scRNA-seq. Cells are colored based on activity scores of luminal progenitor, mature luminal, and basal gene signatures. **(D)** Ternary plot showing the relative activity of luminal progenitor, mature luminal, and basal gene signatures across all tumor cells, colored by unsupervised cluster. **(E)** Ternary plots showing the distribution of relative activity of luminal progenitor, mature luminal, and basal gene signatures across all tumor cells profiled by scRNA-seq. In the left plot, cells assigned to cluster 15 are highlighted in red, while cells assigned to cluster 16 are highlighted in the right plot. **(F-G)** Representative images of periostin **(F)** and osteopontin **(G)** IHC staining in luminal B and low-ER TMAs. **(H)** Quantified osteopontin expression in luminal B and low-ER TMAs.
